# MCMICRO: A scalable, modular image-processing pipeline for multiplexed tissue imaging

**DOI:** 10.1101/2021.03.15.435473

**Authors:** Denis Schapiro, Artem Sokolov, Clarence Yapp, Jeremy L. Muhlich, Joshua Hess, Jia-Ren Lin, Yu-An Chen, Maulik K. Nariya, Gregory J. Baker, Juha Ruokonen, Zoltan Maliga, Connor A. Jacobson, Samouil L. Farhi, Domenic Abbondanza, Eliot T. McKinley, Courtney Betts, Aviv Regev, Robert J. Coffey, Lisa M. Coussens, Sandro Santagata, Peter K. Sorger

## Abstract

Highly multiplexed tissue imaging makes molecular analysis of single cells possible in a preserved spatial context. However, reproducible analysis of the underlying data poses a substantial computational challenge. Here we describe a modular and open-source computational pipeline (MCMICRO) for performing the sequential steps needed to transform large, multi-channel whole slide images into single-cell data. We demonstrate use of MCMICRO on images of different tissues and tumors acquired using multiple imaging platforms, thereby providing a solid foundation for the continued development of tissue imaging software.

## MAIN

The recent introduction of highly multiplexed tissue imaging makes it possible to measure the levels and localization of 20-100 antigens at subcellular resolution in a preserved 3D environment (see **Table S1** for references). In a research setting, multiplexed imaging provides new insight into molecular properties of tissues and their spatial organization and, in a clinical setting, it promises to augment traditional histopathological diagnosis of disease with the molecular information needed to guide use of targeted and immuno-therapies^1–4^. Inadequate tools for image processing remain a substantial barrier to the routine use of multiplexed tissue imaging, particularly in the case of whole-slide imaging (WSI), in which specimens as large as 5 cm^2^ are imaged in their entirety. Diagnostic histopathology is based on WSI, and the FDA mandates it for medical applications^5,6^. We have also found that multiplexed WSI is essential for accurately quantifying the mesoscale structures that organize tissues^7^. Whole-slide images can contain one terabyte of data, 10^5^ to 10^6^ cells, and involve resolvable structures with spatial scales from 100 nm to over 1 cm. This represents a substantial challenge for computational image analysis.

A goal common to almost all multiplexed tissue analyses is identifying cell locations, phenotypes and states based on the levels and patterns of expression of protein markers. These are usually detected using antibodies, often in conjunction with stains such as hematoxylin and eosin (H&E). Image-based single-cell analysis is a natural complement to spatial and single-cell transcriptomics^8–10^ but faces four computational challenges: (i) image segmentation, the process of subdividing images into areas comprising single cells, is difficult when normal tissue structures are disrupted, cells are densely crowded, and nuclei have irregular morphologies – as in cancer; (ii) the fundamental units of tissue organization are highly variable, and the essential data types are not well defined; (iii) WSI generates very large files that must be available for human inspection (a 50-plex 4 cm^2^ image collected at 0.3μm lateral resolution comprises over 400 GB of data); (iv) image processing algorithms are simultaneously being developed by many research groups in parallel, using different programming languages (some proprietary, such as MATLAB) without consideration of interoperability. Analogous challenges in genomics have been addressed by developing computational pipelines that streamline multi-step data analyses and can also be scaled up to cloud compute environments (e.g., Cumulus for scRNAseq)^11^. The use of pipelines involving software containers (e.g., Docker^12^) and workflow languages^13^ makes it possible for multiple research groups to contribute to and iteratively improve complex computational tasks. In the case of tissue atlases, such as the Human Tumor Atlas Network (HTAN)^14^, multiple laboratories are faced with a common set of data analysis challenges, a further motivation for a standardized computational framework.

In this paper we describe MCMICRO (Multiple Choice MICROscopy), a scalable, modular, and open source image processing pipeline implemented in the Nextflow language^15^ that leverages Docker/Singularity containers^12,16^. We show that MCMICRO can process multiplexed data acquired using at least six different imaging technologies (**Table S1**) and has attributes not found in existing workflows (**Table S2**). These include the ability to select among competing algorithms at key steps in the analysis and interactive training of machine learning models (this is particularly important for image segmentation). In common with other bioinformatics pipelines, MCMICRO is designed to complement rather than replace conventional desktop and server-deployed tools. A wide variety of algorithms can be incorporated into the MCMICRO pipeline using containers, and the results can be visualized using multiple software environments, including napari, QuPath, OMERO and histoCAT (see **Table S2** for details and references).

To create MCMICRO, we re-implemented as open-source software several algorithms previously available in the proprietary language MATLAB (MCQuant for quantifying marker intensities and computing morphology metrics^17^, and S3segmenter for watershed segmentation^18^, spot detection, and local thresholding). We also containerized several open-source algorithms (BaSiC^19^ and Ilastik^20^), and incorporated three algorithms and associated deep learning models developed in our laboratories (UMAP/UnMicst, Coreograph and ASHLAR) (**Fig. 1A;** module names in red). All algorithms were tuned to manage very large files (~ 500GB/image) and containerized to abstract away language-specific dependencies (**Methods**). Source code, a user guide and other training materials are available via GitHub (https://github.com/labsyspharm/mcmicro).

**Fig. 1:**
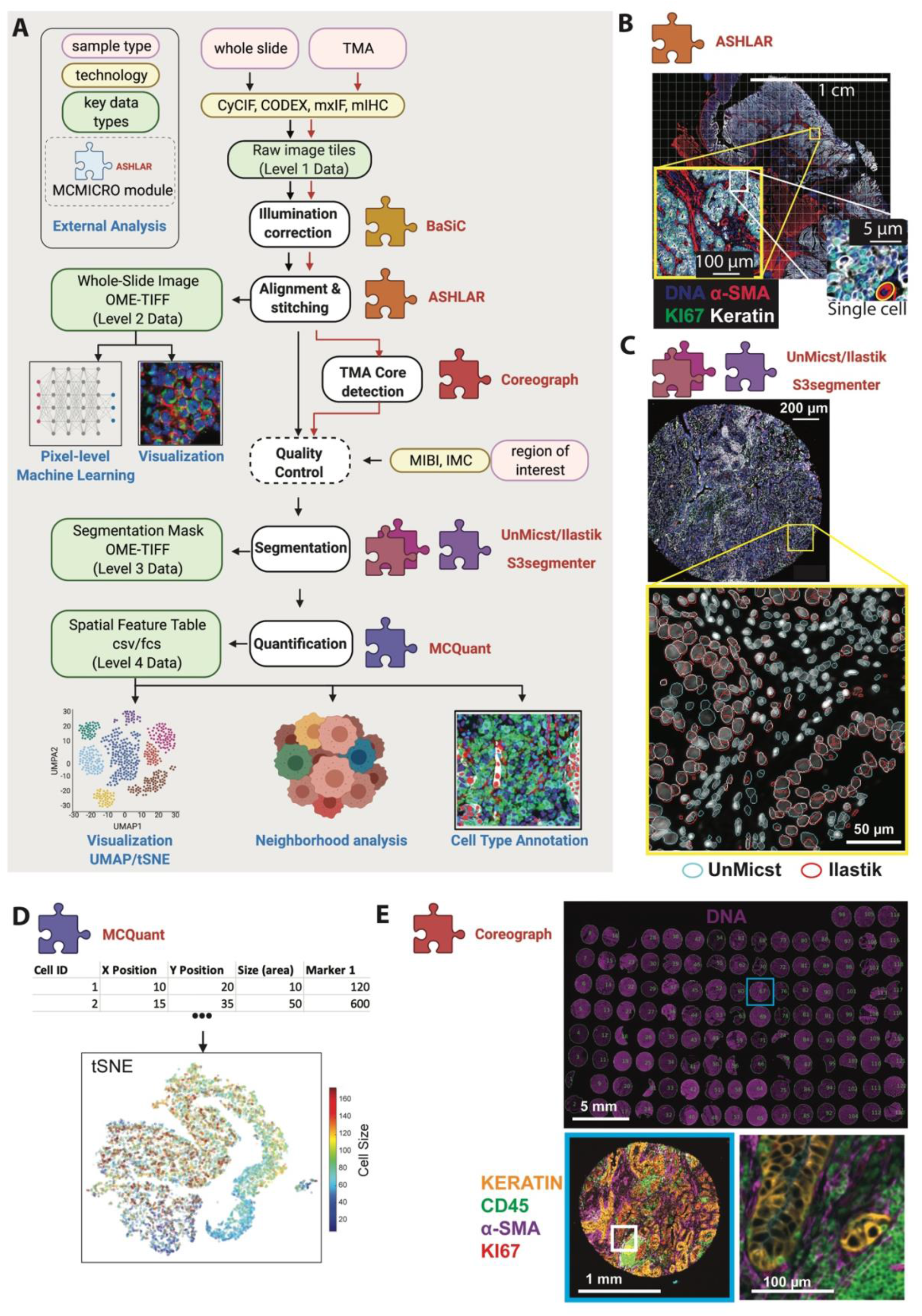
MCMICRO pipeline overview. Modules highlighted in bold red are developed and/or containerized in-house. **(A)** A schematic representation of a canonical workflow for end-to-end image processing of multiplexed whole-slide and TMA using MCMICRO. Shown is a flow of inputs (pink rectangles) from imaging technologies (yellow rectangles) through image processing steps (white rectangles) that are implemented in software modules (puzzle pieces) to produce key data types (green rectangles). Data flows associated with the whole slide and TMA are represented with black and red arrows, respectively. Quality control is highlighted with a dashed border. **(B-E)** Highlights of individual software modules incorporated into MCMICRO. **B** ASHLAR is used to stitch and register individual CyCIF image tiles with subcellular accuracy (yellow zoom-in). This panel depicts a 484 tile (22 x 22) t-CyCIF, whole-slide, mosaic image of a human colorectal cancer in four channels: Hoechst 33342-stained nuclear DNA (blue), α-smooth muscle actin (α-SMA; red), the Ki-67 proliferation marker (green) and cytokeratin (white). An interactive on-line visualization of these data can be found at: https://www.cycif.org/data/tnp-2020/osd-crc-case-1-ffpe-cycif-stack. **C** Two different segmentation masks computed by UnMicst (blue) and Ilastik (red) overlaid on an image of nuclei from an EMIT TMA core. **D** A schematic of the first rows and columns of a Spatial Feature Table used for visualization using tSNE. **E** A CyCIF image of an EMIT TMA de-arrayed using Coreograph to identify individual cores, which are subsequential extracted and analyzed in high-resolution. Below, a five-color image of a single lung adenocarcinoma core is shown for channels corresponding to Hoechst 33342-stained DNA (white), cytokeratin (orange), the immunecell marker CD45 (green), α-SMA (magenta) and Ki-67 (red)).

To facilitate benchmarking, development of new algorithms and model training, we also generated a set of freely available reference images, the Exemplar Microscopy Images of Tissues and Tumors (EMIT). EMIT comprises multiplexed CyCIF images of a tissue microarray (TMA) with 120 1.5 mm cores from 34 types of cancer, non-neoplastic diseases, and matched normal tissue (**Figure S1,** https://synapse.org/EMIT). EMIT images were processed using MCMICRO (using the Coreograph module) and all steps are documented on Synapse (https://synapse.org/EMIT). Clustering of normal tissues and cancers by type (with some variance, because specimens came from different individuals) demonstrates that a wide range of specimens can be processed by MCMICRO to generate meaningful single cell data **(Figure S2).**

Processing multiplexed WSI data starts with acquisition of individual image tiles in a BioFormats-compatible format (level 1 data; **Fig. 1A**)^21^; each tile is typically a megapixel multichannel image, and as many as 10^3^ tiles are required to cover a large tissue specimen at subcellular resolution. Tiles are corrected for uneven illumination, stitched together, and registered across channels to generate the first broadly useful type of data: a fully assembled, multichannel *mosaic image* in OME-TIFF format (a class of level 2 data) (**Fig. 1B**). In a large mosaic whole-slide image, length scales vary 10^5^-fold from the smallest resolvable feature to the largest dimension. Images are subjected to quality control, followed by segmentation. A segmentation mask (level 3 data), the next object computed by MCMICRO, is available for human inspection in conjunction with underlying images to determine the quality of different segmentation approaches (**Fig. 1C**).

Following segmentation, the staining intensity in each channel is computed on a per-cell basis to generate a *Spatial Feature Table* (level 4 data), which is analogous to a count table in scRNAseq. In its simplest form, this table consists of the positions of cells and their integrated staining intensities in each imaging channel (morphological data, such as size, eccentricity etc. are additional table elements; **Fig. 1D**). The Spatial Feature Table can be visualized using tools designed for high dimensional data such as tSNE or UMAP, processed to identify cell types, and used for neighborhood or other types of analysis (**Fig. 1D**). It is also possible to skip segmentation altogether and perform analysis directly on images; pixel-level deep learning has already shown promise in clinical settings^22,23^, and many algorithms have been generalized for use with multiplexed data. Regardless of how data flows through MCMICRO, provenance is maintained by recording the identities, version numbers and parameter settings for each module, enabling full reproducibility (**Fig. S3**).

MCMICRO includes a newly developed tool for processing TMAs, which are widely used in research, because they enable parallel analysis of many specimens. In a TMA, a single slide carries dozens to hundreds of 0.3 to 2 mm diameter “cores”. The *Coreograph* module in MCMICRO is based on the U-Net deep learning architecture^24^. It finds the locations of individual cores and extracts each core as a separate, multi-channel image (**Fig. 1E**), allowing all cores to be processed in parallel by downstream modules. The robustness of the underlying neural network makes it possible for the module to accurately identify cores even in highly distorted TMAs.

Image processing requires user interaction and frequent visual review (see CellProfiler, for example^25^). To enable human-in-the-loop analysis, MCMICRO allows for training and parameter adjustment to take place locally, using subsets of a large mosaic image. This iterative approach is particularly important for segmentation, since most contemporary algorithms rely on supervised machine learning. An absence of well-defined objective functions and ground truth data makes automated scoring of algorithms difficult, and different combinations of algorithms and models may be optimal for different tissues. MCMICRO therefore incorporates multiple segmentation algorithms (e.g., U-Net^24^ or ilastik^20^), which can be executed in parallel and then compared **(Fig. 1C)**. Additional improvement in segmentation can be achieved with the help of the EMIT data repository (https://synapse.org/EMIT), which includes a “classifier zoo” comprising a set of tissue-specific random forest segmentation models for ilastik. These models aid generation of robust tissue-specific segmentation masks and can also be subjected to further dataset-specific training.

To demonstrate the technology-agnostic capabilities of MCMICRO, we collected data from a single FFPE tonsil specimen at four different institutions using four imaging technologies: CODEX and CyCIF, which are immunofluorescence-based; mIHC, which uses multiplexed immunohistochemistry; and H&E staining **(Fig. 2A)**. We also analyzed mxIF and publicly available Imaging Mass Cytometry (IMC) and MIBI data (**Table S2**). To show that MCMICRO does not have specific hardware dependencies, data processing was performed using cloud compute nodes provided either by Amazon Web Services (AWS) or the Google Cloud Platform and also using a Linux-based institutional cluster running the SLURM workload manager. MCMICRO provides detailed information on time, memory and CPU usage, making it straightforward to provision necessary computational resources (**Fig. S4**).

**Fig. 2:**
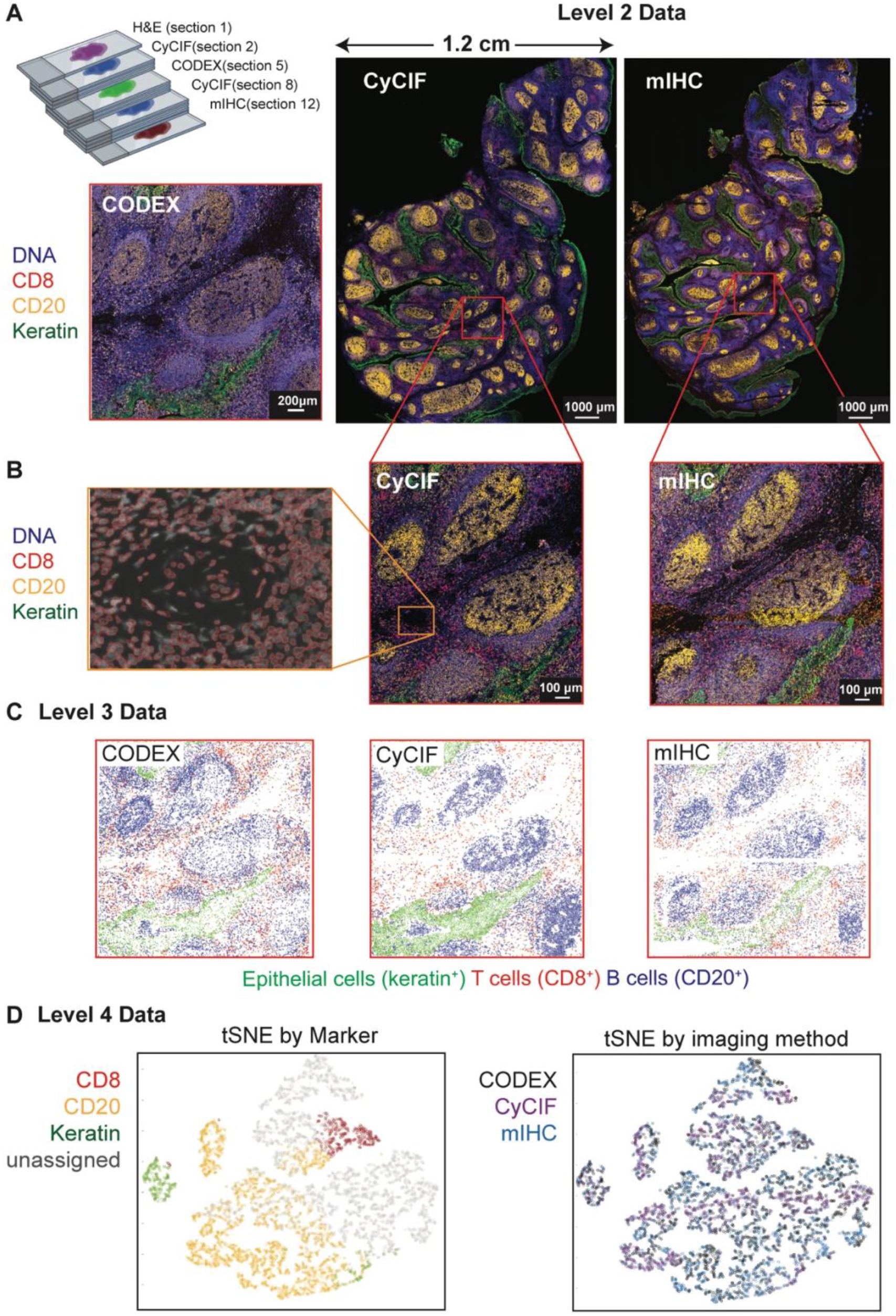
Comparison of images of human tonsil collected using three different technologies and processed using MCMICRO. **A.** Serial sections of a single tissue block imaged using H&E, 11-marker CODEX, 27-marker CyCIF, and 16-marker mIHC. The sectioning plan shows the position of each 5 μm section within the block: H&E section 1, CyCIF section 2, CODEX section 5 and mIHC section 12. Images show selected channels as follows: Hoechst 33342 (blue), CD20 (orange), Keratin (green), and CD8 (red). The CODEX image shows only a specific region (red border) of the specimen visible in whole-slide images to the right. **B.** Higher magnification images of the data above highlighting individual cells and segmentation masks generated with UnMicst. **C.** Centroids of the single cell mask for CODEX, CyCIF and mIHC are colored by marker expression to identify cell types. Epithelial cells of the tonsil mucosa stain positive for pan-cytokeratin (green), cytotoxic T cells stain positive for CD8 (red); and B cells stain positive for CD20 (blue). **D.** t-SNE of combined CODEX, CyCIF and mIHC data demonstrating clustering by marker expression (left) but not imaging technology (right).

Image tiles from a variety of microscopes were subjected to stitching, registration and illumination correction using ASHLAR and BaSiC to generate mosaic level 2 image data that was visually inspected on a local workstation using napari and in the cloud using OMERO **(Fig. 2A)**. Images were then segmented and staining intensities were computed on a per-cell basis using MCQuant. Cell types were visualized in the tissue context for epithelial cells of the tonsil mucosa (Keratin+/panCK+), cytotoxic T cells (CD8+) and B cells (CD20+) **(Fig. 2B)**. Visual inspection of stitched and registered CyCIF, CODEX and mIHC images and derived data revealed accurate image stitching and registration, facilitating the creation of reasonable segmentation masks and the generation of correctly formatted Spatial Feature Tables. The results of cell type calling were similar **(Fig. 2C),** and when data from all three technologies were combined and visualized using tSNE, cells were separated by marker expression not imaging technology **(Fig 2D).** These findings demonstrate consistency in image acquisition and data processing.

A few algorithms in MCMICRO (e.g., Ashlar and BaSIC) are tissue and technology agnostic and can be used on diverse types of data with little, if any, tuning or modification. The performance of other algorithms (e.g., UnMicst and Ilastik) is dependent on the properties of their learned models, which often work well for some tissues and not for others. MCMICRO facilitates identification of effective algorithms and models by executing different segmentation approaches in parallel, followed by comparison of the resulting masks. We expect continued innovation in the area of image segmentation, as well as addition of algorithms for automated quality control of images and identification of cell types based on marker intensities and cell morphologies. However, we do not anticipate that users will need to manage an endless proliferation of novel methods: multiple research consortia are actively working together on evaluation efforts (analogous to Dream Challenges^26^) aimed at creating best practices for highly-multiplexed image analysis. MCMICRO provides the technical foundation for such evaluations and for widespread distribution of the results. MCMICRO is also being used by the HTAN consortium to rigorously compare different image acquisition technologies.

In conclusion, the MCMICRO pipeline described here provides a foundation for community-wide development of FAIR (findable, accessible, interoperable and reusable)^27^ workflows for analysis of large tissue images currently being generated by multiple international consortia and many individual laboratories. MCMICRO works with any acquisition technology that generates Bio-Formats/OME-compatible images, including the six technologies described above. The pipeline is based on widely accepted software standards and interoperates with any programming language through the use of software containers, making it easy to add new modules. The result is a user-friendly end-to-end pipeline that executes computation-intensive processes in the cloud, while enabling parameter optimization, training and quality control to be performed locally and interactively.

## METHODS

### Tissue samples

A de-identified tonsil specimen from a 4-year old Caucasian female was procured from the Cooperative Human Tissue Network (CHTN), Western Division, as part of the Human Tumor Atlas (HTAN) SARDANA trans-network project (TNP). Regulatory documents including Institutional Review Board (IRB) protocols, data use agreements and tissue use agreements were in place to ensure regulatory compliance. Standard protocols for tissue procurement and fixation were followed; a detailed protocol can be found at the link provided in Table 1. Sections were cut from a common formalin-fixed paraffin embedded (FFPE) block at a thickness of 5 μm and mounted onto Superfrost Plus glass microscope slides (Fisher Scientific, 12-550-15) for CyCIF and mIHC or mounted on poly-L-Lysine (PLL) coated coverslips (Electron Microscopy Sciences, 72204-01; slides and FFPE sections prepared following instructions in the Akoya Biosciences CODEX User Manual Rev B.0, Chapter 3. Coverslip Preparation and Tissue Processing) for CODEX. A set of FFPE tissue sections was received by participating HTAN Centers, as indicated in **Table 1**, allowing Centers to generate a comparable spatial cell census using each Center’s imaging method of choice. CHTN performed hematoxylin and eosin (H&E) staining on the first section which was subsequently imaged at Harvard Medical School (HMS).

**Table 1.** Sample information.

| Section Number | Section Thickness ($\mu\text{m}$) | Center | Assay |
| --- | --- | --- | --- |
| WD-75684-01 | 5 | Cooperative Human Tissue Network | H&E |
| WD-75684-02 | 5 | Harvard Medical School | CyCIF |
| WD-75684-05 | 5 | Broad Institute | CODEX |
| WD-75684-08 | 5 | Harvard Medical School | CyCIF |
| WD-75684-12 | 5 | Oregon Health & Science University | mIHC |

For the EMIT dataset, human tissue specimens (from 42 patients) were used to construct a multi-tissue microarray (HTMA427) under an excess (discarded) tissue protocol approved by the IRB at Brigham and Women’s Hospital (BWH IRB 2018P001627). Two 1.5 mm diameter cores were acquired from each of 60 tissue regions with the goal of acquiring one or two examples of as many tumors as possible (with matched normal tissue from the same resection when that was feasible), as a well several non-neoplastic medical diseases involving acute inflammation (e.g. diverticulitis and appendicitis), and secondary lymphoid tissues such as tonsil, spleen and lymph nodes. Overall, the TMA contained 120 cores plus 3 additional “marker cores,” which are cores added to the TMA in a manner that makes it possible to orient the TMA in images.

### CyCIF staining and imaging

The CyCIF method involves iterative cycles of antibody incubation, imaging and fluorophore inactivation as described previously^7^. A detailed protocol can be found on protocols.io as shown in **Table 2**. CyCIF images are 36-plex whole-slide images collected using a 20x magnification, 0.75 NA objective with 2 x 2 pixel binning, yielding images of pixel size at 0.65 μm/pixel. The image comprises 416 and 350 image tiles for WD-75684-02 and WD-75684-08, respectively, each with four channels, one of which is always Hoechst to stain DNA in the nucleus.

**Table 2.**
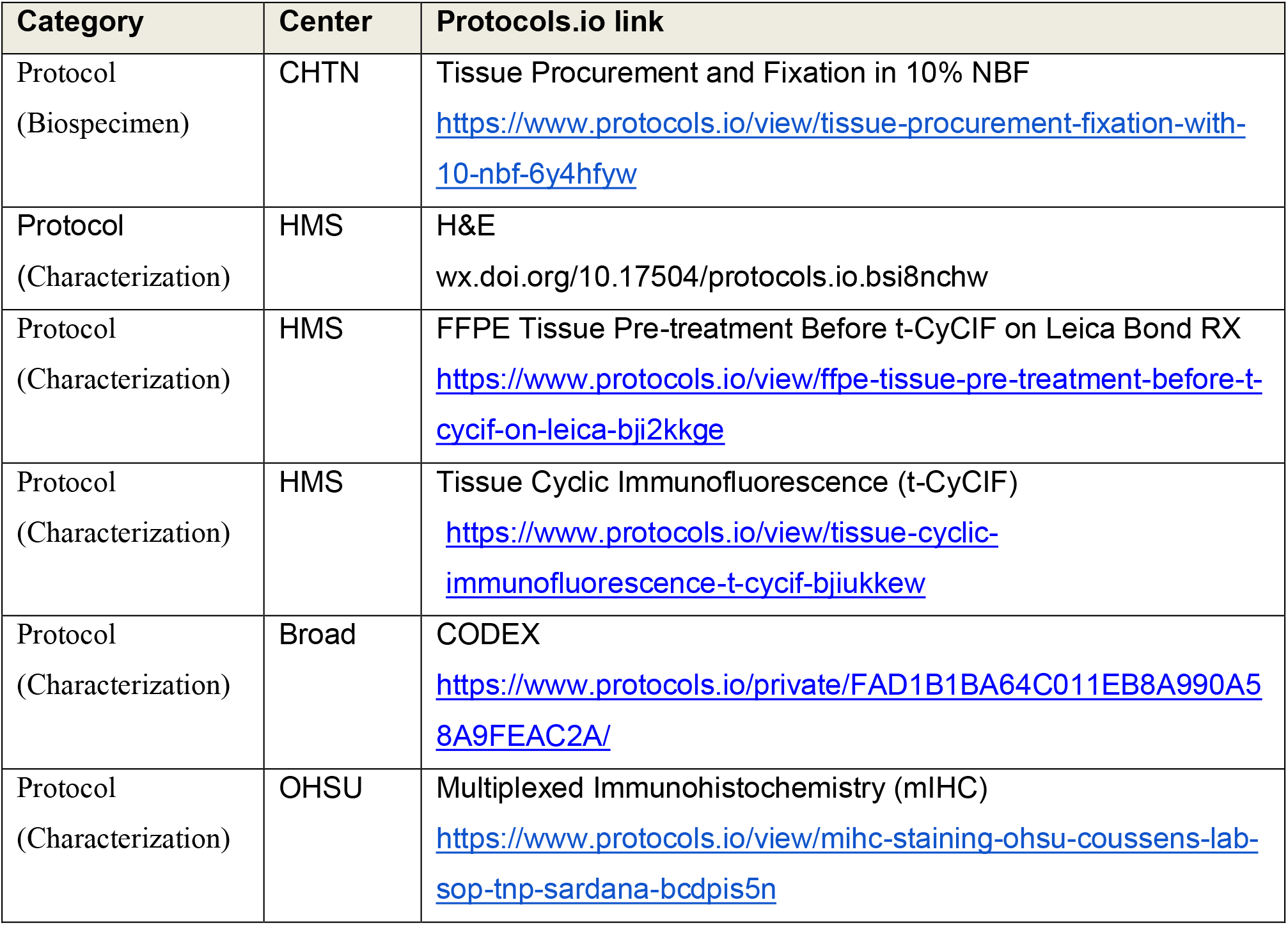
List of protocols. As a part of the HTAN effort, all protocols and methods are deposited on Protocols.io.

### CODEX staining and imaging

Coverslips were prepared following the FFPE tissue staining protocols in the Akoya Biosciences CODEX User Manual (Sections 5.4 – 5.6). Briefly, 5 μm FFPE tissue sections were cut onto PLL-coated coverslips and baked for 20-25 minutes at 55 °C. Sections were cooled briefly before deparaffinization and were washed for 5 minutes each as follows: twice in xylene, twice in 100% ethanol, once in 90%, 70%, 50%, and 30% ethanol, and twice in deionized water. Sections were moved to 1x Citrate Buffer (Vector Laboratories, H-3300) and antigen retrieval was performed in a Tinto Retriever Pressure Cooker (BioSB, BSB 7008) at high pressure for 20 minutes. Sections were briefly washed in deionized water before being left to incubate in deionized water at room temperature for 10 minutes. Sections were briefly washed twice in Hydration Buffer (Akoya), then were left to incubate in Staining Buffer (Akoya) at room temperature for 20-30 minutes. 200 μL/section of Antibody Cocktail was prepared according to manufacturer instructions. Sections were covered with the 200 μL Antibody Cocktail and left to incubate at room temperature for 3 hours in a humidity chamber. Sections were washed twice in Staining Buffer for 2 minutes, and then fixed with a mixture of 1.6% PFA in Storage Buffer (Akoya) for 10 minutes. Sections were briefly washed three times in 1x PBS, and then washed in ice-cold methanol for 5 minutes before being washed again three times in 1x PBS. Sections were stained with 190 μL of a mixture of 20 μL Fixative Reagent (Akoya) and 1 mL 1x PBS, after which they were left to incubate at room temperature for 20 minutes. Sections were briefly washed three times in 1X PBS and were stored in Storage Buffer at 4 °C until the assay was ready to be run.

### Running the CODEX Assay

A 96-well plate of reporter stains with Nuclear Stain (Akoya) was prepared according to Akoya Biosciences CODEX User Manual (Sections 7.1 – 7.2). Stained Tissue sections were loaded onto the CODEX Stage Insert (Akoya) and the Reporter Plate was loaded into the CODEX Machine. The on-screen prompts were followed and the section was manually stained with a 1:2000 Nuclear Stain in 1x CODEX Buffer (Akoya) for 5 minutes before proceeding with following the on-screen prompts. Imaging was performed on a Zeiss Axio Observer with Colibri 7 light source. Emission filters were BP 450/40, BP 550/100, BP 525/50, BP 630/75, BP 647/70, BP 690/50, and TBP 425/29 + 514/31 + 632/100 and dichroic mirrors were QBS 405 + 492 + 575 + 653, TFT 450 + 520 + 605, TFT 395 + 495 + 610, and TBS 405 + 493 + 575, all from Zeiss. Overview scans were performed at 10x magnification, after which 5 x 5 field of view regions were acquired using a Plan-Apochromat 20x/0.8 M27 Air objective (Zeiss, 420650-9902-000). 20x magnification images were acquired with a 212 x 212 nm pixel size using software autofocus repeated every tile before acquiring a 17 plane z-stack with 0.49 μm spacing. Tiles were stitched using a 10% overlap.

### mIHC staining and imaging

The multiplex immunohistochemistry (mIHC) platform described herein involves wet and dry-lab techniques that have been robustly developed to interrogate the tumor immune microenvironment in situ. mIHC involves a cyclic staining process optimized for FFPE tissues with panels of antibodies (12-29 per panel) designed to interrogate both lymphoid and myeloid compartments of the immune system as well as cellular functional states, as previously described^28,29^.

### Pipeline implementation

MCMICRO was implemented in Nextflow, which was chosen for its natural integration with container technologies such as Docker and Singularity, its automatic provenance tracking and parallelization of image processing tasks, and its ability to specify module dependencies that may change at runtime^15^.

### Illumination correction

BaSiC is a Fiji / ImageJ plugin for background and shading correction, producing high accuracy while requiring only a few input images^19^. We containerized the tool, allowing it to be executed without an explicit installation of ImageJ.

### Image stitching and registration using Ashlar

Cycle-based highly multiplexed microscopy produces multi-channel images of fixed cells using a standard four/five-color microscope. Registration of the images across successive cycles is made straightforward by the addition of a nuclear counterstain in every cycle. Given a set of slightly overlapping images covering a tissue, we correct for mechanical stage positioning error intrinsic to all microscopes using Ashlar (Alignment by Simultaneous Harmonization of Layer/Adjacency Registration), a Python package for efficient mosaicing and registration of highly multiplexed imagery^30^. The overall strategy of Ashlar is as follows: (i) align tile images from the first cycle edge-to-edge with their nearest neighbors (mosaicing) using phase correlation on the nuclear marker channel; (ii) for the second and subsequent cycles, align each tile to the greatest overlapping tile from the first cycle (registration), using phase correlation on the nuclear marker channel, and retain the corrected stage coordinates, rather than the actual merged images; (iii) use the corrected coordinates to assemble a single image covering the entire imaged area, including all channels from all cycles. This approach minimizes the compounding of alignment errors across tiles and cycles as well as temporary storage requirements for intermediate results.

### Coreograph

Coreograph’s function is to split, or ‘dearray’, a stitched TMA image into separate image stacks per core. It employs a semantic segmentation preprocessing step to assist with identifying cores that are dimmed or fragmented, which is a common issue. We trained a deep, fully connected network on two classes – core tissue and background – using the popular UNet^24^ architecture for semantic segmentation. Training data consisted of cores that were well-separated, as well as cores that were merged and/or fragmented, which allowed for handling situations where sample integrity was highly heterogeneous. Once cores had been accentuated in the form of probability maps, they were cropped from the stitched image based on their median diameter and saved as a TIFF stack. In situations where the cores were too clumped, the median diameter was used to set the size of a Laplacian of Gaussian (LoG) kernel in order to identify local maxima from the probability maps.

### UnMicst (U-Net model for identifying cells and segmenting tissue)

UnMicst is a preprocessing module in MCMICRO that aids in improving downstream segmentation accuracy by generating per-class probability maps to classify each pixel with a certain amount of confidence. Analogous to Coreograph, it employs a UNet architecture (see above). Previously, a similar UNet model was trained for nuclei segmentation to recognize two classes in Hoechst 33342 – stained tonsil tissue (nuclei contours and background). Here, we train a 3-class model to extract nuclei centers, nuclei contours, and background from manually annotated lung, tonsil, prostate and other tissues in order to ascribe a variety of nuclei shapes. Realistic augmentations, in addition to conventional on-the-fly transformations, were included by deliberately defocusing the image and increasing the exposure time of the camera to simulate focus and contrast augmentations, respectively. Training was performed using a batch size of 24 with the Adam Optimizer and a learning rate of 0.00003 until the accuracy converged. Segmentation accuracy was estimated by counting the fraction of cells in a held out test set that passed a sweeping Intersection of Union (IOU) threshold.

### Ilastik tissue segmentation

Similar to UnMicst, Ilastik assigns each pixel a probability of belonging to predetermined classes (e.g., cell nucleus, membrane, background). MCMICRO relies on Ilastik’s pixel classification module for training and subsequent batch-processing using a random forest classifier. Ilastik classifier training in MCMICRO is completed in several steps. First, regions of interest (ROIs) with a user-defined width and height are randomly cropped from the WSI. Second, the ROIs are manually annotated by the user on a local machine via Ilastik’s graphical user interface (GUI). Third, to ensure tissue portions are accurately represented in cropped images, Otsu’s method is used to identify a global threshold across the WSI for a particular channel of interest (e.g., nuclear staining). Finally, the user exports the cropped sections that contain the desired proportion of pixels above the previously determined threshold. Upon completion of the random forest training, whole slide classifier predictions are deployed in headless mode (no GUI) for batch processing of large data sets within MCMICRO.

### Watershed segmentation via S3segmenter

We implemented S3segmenter, a custom marker-controlled watershed algorithm to identify nuclei from the probability maps generated by UnMicst and Ilastik. Watershed markers are obtained by convolving a LoG kernel, followed by a local maxima search across the image to identify seed points. The size of the LoG kernel and local maxima compression are tunable parameters dependent on the expected nuclei diameters in the image. As a byproduct, this method identifies false positive segments in the image background. These false positives were excluded by comparing their intensities to an Otsu-derived threshold calculated either on the raw image or on the probability map. S3segmenter currently offers three alternative methods for cytoplasm segmentation. First, traditional nonoverlapping rings (annuli) with user-defined radius are used around each nucleus. Second, a Euclidean distance transform is computed around each nucleus and masked with a user-specified channel, reflecting the overall shape of the whole tissue sample. An autofluorescence channel can be chosen if the signal-to-image background ratio is sufficiently high. Third, the cytoplasm is segmented using a marker-controlled watershed on the grayscale-weighted distance transform, where the segmented nuclei are markers and the grayscale-weighted distance transform is approximated by adding scaled versions of the distance transform and raw image together. This method is conceptually similar to that found in the CellProfiler Identify Secondary Objects module^25^. S3segmenter is also capable of detecting puncta by convolving a small LoG kernel across the image and identifying local maxima. Once nuclei and cytoplasm segmentation are complete, labelled masks for each region are exported as 32-bit tiff images. Two channel tiff stacks consisting of the mask outlines and raw image are also saved so that segmentation accuracy can be easily visually assessed.

### MCQuant

Semantic segmentation in MCMICRO produces 32-bit masks, which are used to quantify pixel intensity (i.e., protein expression) on multiplexed WSI for cytoplasm and nuclei. Quantification in MCMICRO is carried out using scikit-image, a popular Python-based image analysis library, and values of cellular spatial features are calculated for unique cells (cytoplasm and nuclei), in addition to their mean pixel intensity (protein expression). The resulting spatial feature tables are exported as CSV files for subsequent data analysis analogous to histoCAT^17^, which is implemented in MATLAB.

### Data availability statement

All software and code that produced the findings of the study, including all main and supplemental figures, are available at https://github.com/labsyspharm/mcmicro.

All EMIT images are available at https://synapse.org/EMIT and all exemplar and tonsil images are available at https://synapse.org/MCMICRO_images.

## ACKNOWLEDGEMENTS

This work was funded by NIH grants U54-CA225088 and U2C-CA233262 to P.K.S. and S.S and by the Ludwig Cancer Center at Harvard. D.S. was funded by an Early Postdoc Mobility fellowship (no. P2ZHP3_181475) from the Swiss National Science Foundation and is a Damon Runyon Fellow supported by the Damon Runyon Cancer Research Foundation (DRQ-03-20). Z.M. is supported by NCI grant R50-CA252138. We thank Dana-Farber/Harvard Cancer Center in Boston, MA, for the use of the Specialized Histopathology Core, which provided TMA construction and sectioning services. Dana-Farber/Harvard Cancer Center is supported in part by an NCI Cancer Center Support Grant P30 CA06516. Tissue samples were provided by the NCI Cooperative Human Tissue Network (CHTN).

## OUTSIDE INTERESTS

PKS is a member of the SAB or BOD member of Applied Biomath, RareCyte Inc., and Glencoe Software, which distributes a commercial version of the OMERO database; PKS is also a member of the NanoString SAB. In the last five years the Sorger lab has received research funding from Novartis and Merck. Sorger declares that none of these relationships have influenced the content of this manuscript. SS is a consultant for RareCyte Inc. A. R. is a cofounder and equity holder of Celsius Therapeutics, an equity holder in Immunitas and, until 31 July 2020, was an SAB member of Thermo Fisher Scientific, Syros Pharmaceuticals, Neogene Therapeutics and Asimov. Since August 1, 2020, A. R. has been an employee of Genentech. The other authors declare no outside interests.

---

**Figure S1.**
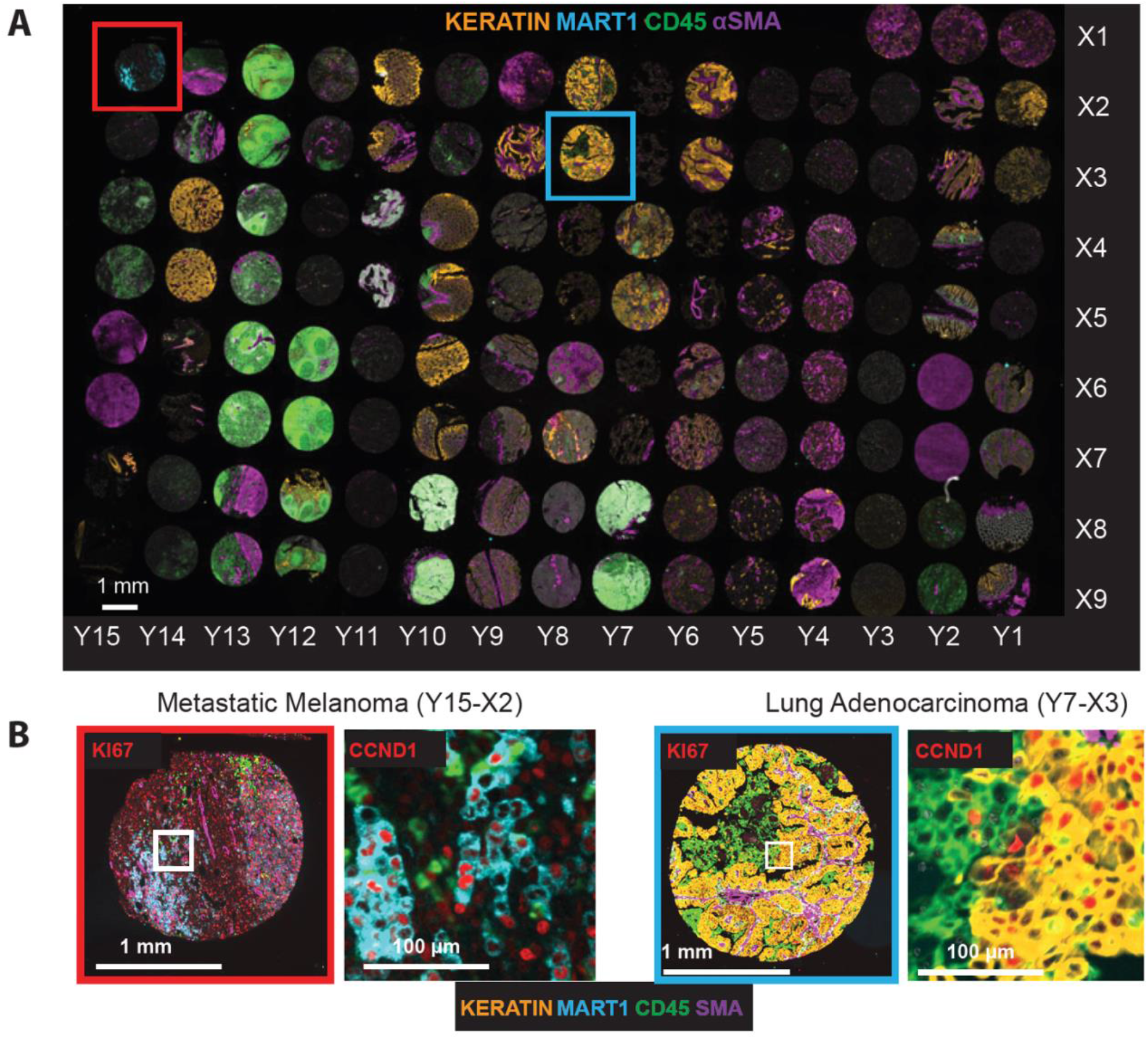
The EMIT dataset spanning 123 tissue cores across 34 cancer, non-neoplastic diseases, and normal tissue type. **A.** CyCIF whole slide image of EMIT visualizing Hoechst 33342-stained nuclear DNA (white), Keratin (orange), MART1 (cyan), CD45 (green) and SMA (purple). **B. A** zoom-in view of a metastatic melanoma (left, red box) and a lung adenocarcinoma (right, blue box) core. The highest zoom level is highlighted with white boxes in the corresponding low magnification images.

**Figure S2.**
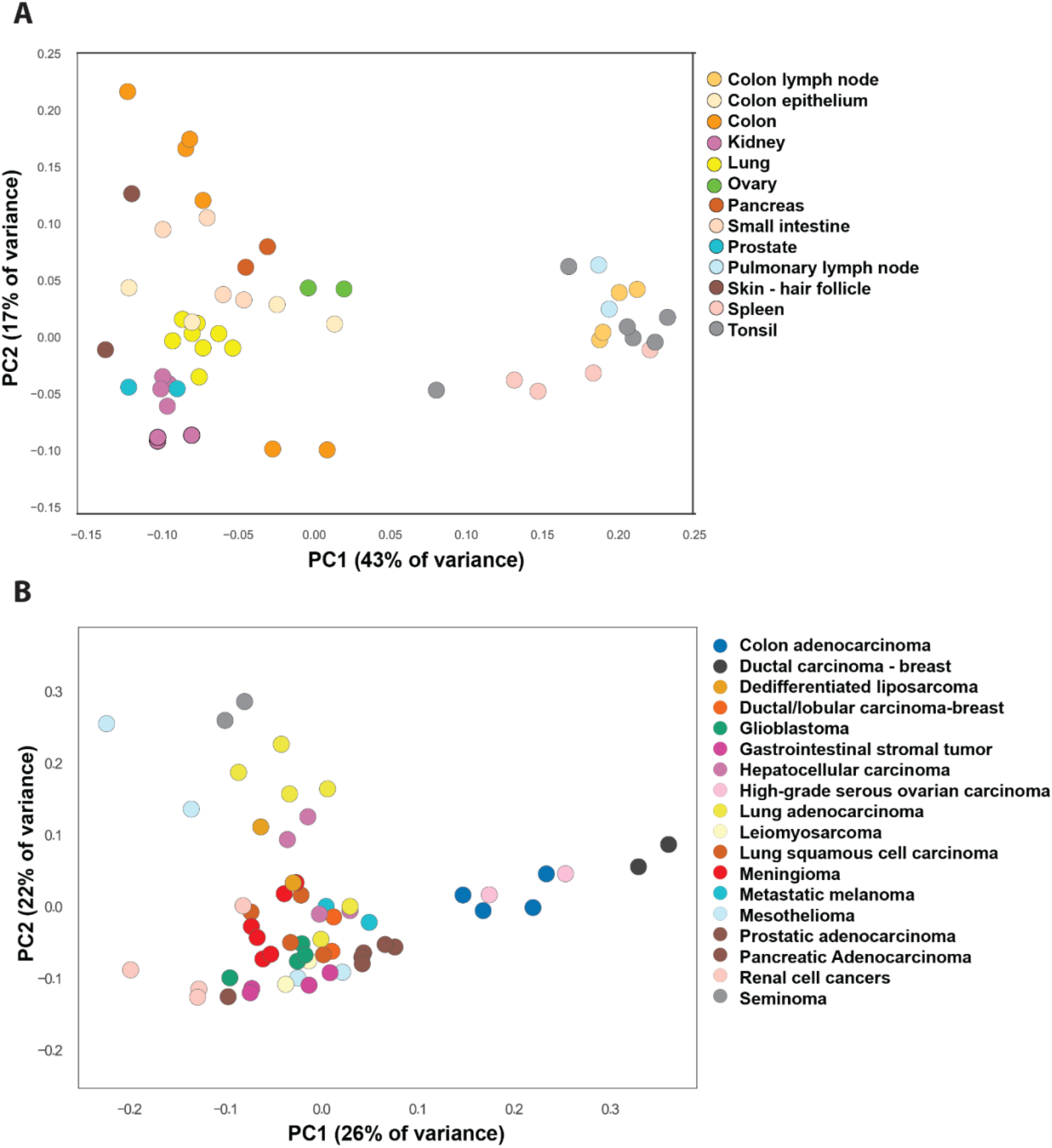
Principal component analysis (PCA) of Spatial Feature Tables derived from EMIT images. **A.** represents normal tissues and **B.** cancer tissues. Independent cores cluster to a substantial degree by tissue or cancer type; some variation is expected because tumors had different grades and derive from different individuals. Data from the following antibodies was used to generate the data: CD73, MART1, KI67, pan-cytokeratin, CD45, ECAD, α-SMA, CD32, CDKN1A, CCNA2, CDKN1C, CDKN1B, CCND1, cPARP, CCNB1, PCNA and CDK2.

**Figure S3.**
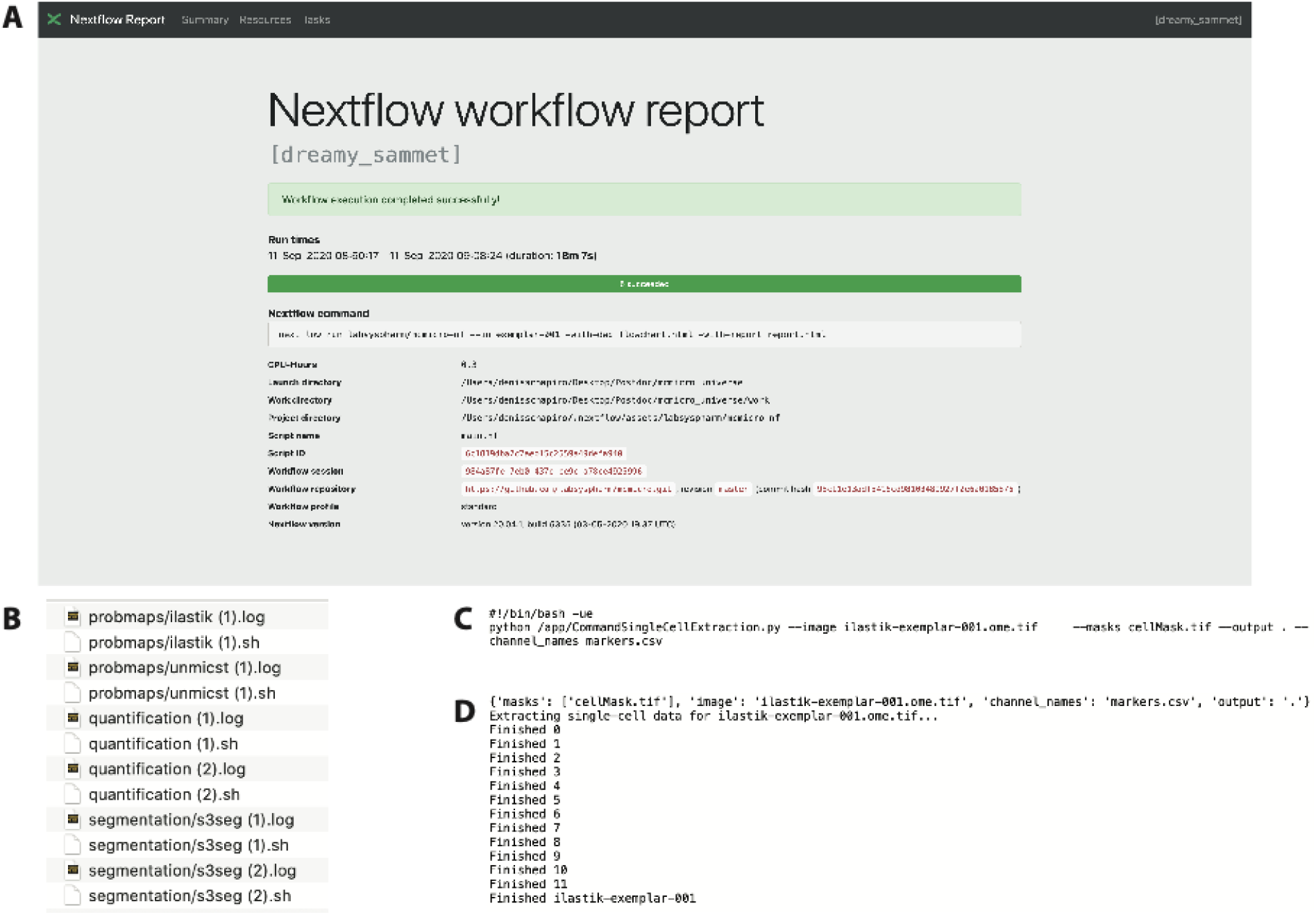
Nextflow enables reproducible data processing using the provenance module. **A.** Nextflow report provides detailed documentation for used resources, directories, repositories (including commit hash) and the corresponding execution times. The report is browser based and interactive. **B-D.** Provenance reconstruction enabled by recording each executed command (.sh) and its output (.log). Representative examples of a command and its output are shown in (C) and (D), respectively.

**Figure S4.**
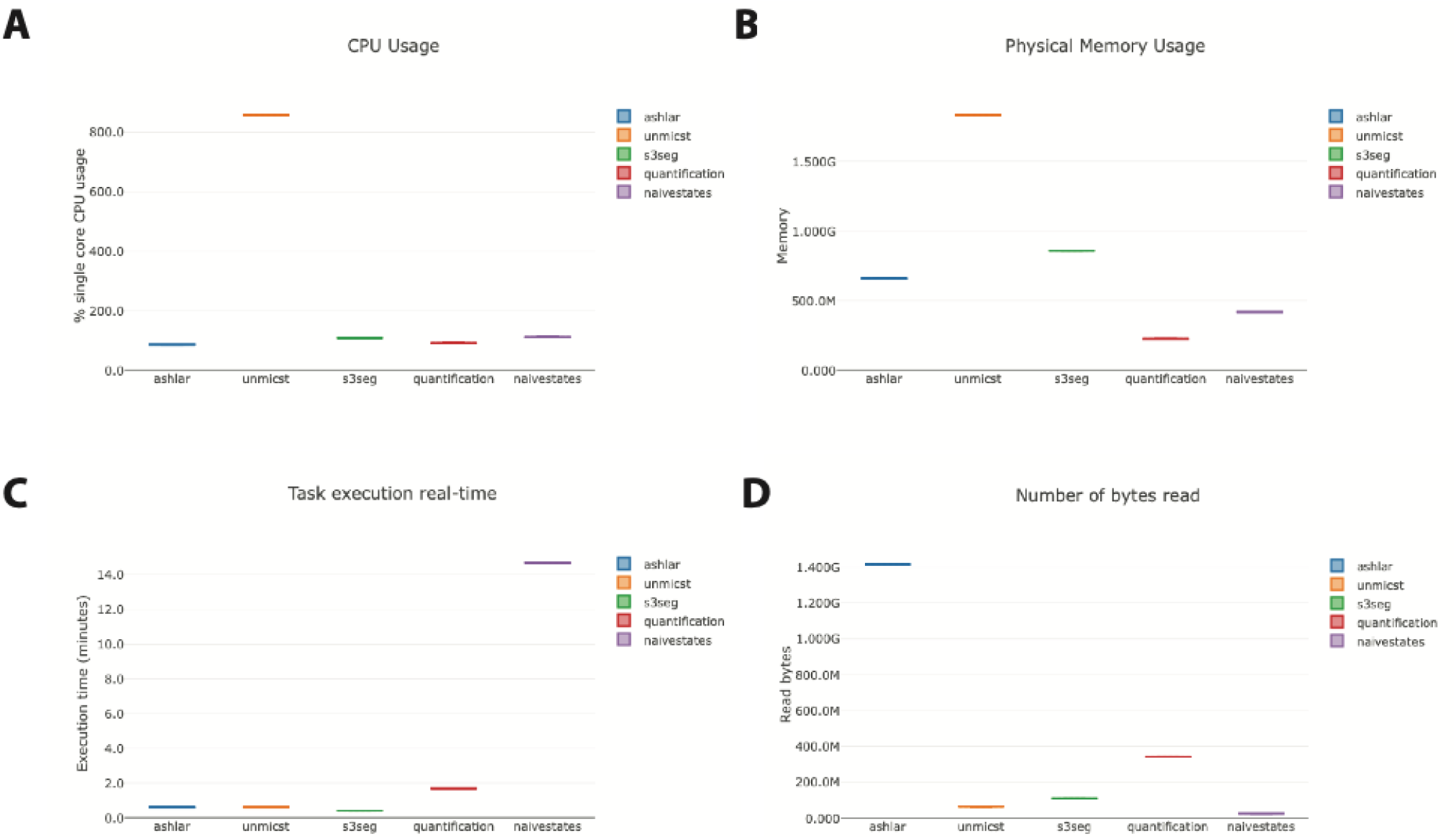
Detailed insight into the computational resources required by each module, generated by Nextflow. The data is viewed as an interactive browser-based report. **A.** CPU usage is recorded as either % single core CPU usage (visualized) or % CPUs allocated. **B.** Physical memory usage is recorded as either RAM only (visualized), RAM + Disk swap or % RAM allocated. **C.** Job duration is recorded as either execution time (visualized) or % time allocated. **D.** Input/Output (I/O) records both read (visualized) and written bytes.

**Table S1:** Highly multiplexed imaging methods. Orange labeled methods were successfully processed by MCMICRO on publicly available datasets. Green labeled methods are additionally tested on images unique to this study with detailed description in the documentation.

| Non-cyclic metal-based | Cyclic fluorescence imaging |  | Cyclic immune-histochemistry |
| --- | --- | --- | --- |
|  | cyclic-stained | single-stained |  |
| Imaging Mass Cytometry (IMC)<sup>1</sup> | CODEX<sup>2</sup> | MxIF<sup>3</sup> | mIHC<sup>4</sup> |
| Multiplexed Ion Beam Imaging (MIBI)<sup>5</sup> | Immuno-SABER<sup>6</sup> | CyCIF<sup>7</sup> | MICSSS<sup>8</sup> |
|  |  | 4i<sup>9</sup> |  |

**Table S2:** Available open-source tools for image processing, analysis and visualization. List of open-source tools available for highly multiplexed image processing.

| Software | Scalable | Whole slide processing | Stitching and registration | Modular | Segmentation | Analysis | GUI |
| --- | --- | --- | --- | --- | --- | --- | --- |
| MCMICRO | Yes | Yes | Yes | Yes | Yes | Yes | No |
| Cytokit<sup>10</sup> | Yes | No (tiles) | No | No | Yes | Yes | Yes |
| starfish<sup>11</sup> | Yes | No (tiles) | No | Yes | Yes | Yes | No |
| histoCAT<sup>12</sup> | No | Yes | No | Yes | No | Yes | Yes |
| QuPath<sup>13</sup> | No | Yes | No | No | Yes | Yes | Yes |
| CytoMAP<sup>14</sup> | No | Yes | No | No | No | Yes | Yes |
| Facetto<sup>15</sup> | No | Yes | No | No | No | Yes | Yes |
| napari<sup>16</sup> | Visualization tool |  |  |  |  |  |  |
| OMERO<sup>17</sup> | Visualization tool |  |  |  |  |  |  |
| Minerva<sup>18</sup> | Visualization tool |  |  |  |  |  |  |

